# Data-Driven Feed Optimization for Sustainable Aquaculture through Integrated Omics Analysis and Two-stage Bayesian optimization

**DOI:** 10.1101/2025.01.27.635187

**Authors:** Kei Terayama, Satoshi Soma, Hirofumi Furuita, Tomohito Shimizu, Takashi Iwasaki, Kenji Sakata, Ken-ichi Akagi, Taiga Asakura, Tetsuya Sanda, Tomofumi Yamaguchi, Yuji Fujikura, Yuki Hongo, Hideaki Shima, Hideo Yokota, Jun Kikuchi, Motoshige Yasuike, Miyuki Mekuchi

## Abstract

In aquaculture and livestock farming, developing feed that promotes efficient growth while minimizing environmental impact is crucial for sustainability. We developed a data-driven feed optimization system (DFOS) that integrates omics analysis with machine learning-based optimization technologies. First, DFOS employed Bayesian optimization (BO) to identify the optimal proportions of proteins and lipids, which are crucial feed components. Simultaneously, omics analysis was conducted to assess the physiological response of the targets to a given feed and identify key dietary components essential for growth. Next, BO was used to efficiently modify the combination and proportion of the specific additives identified in the first stage. We demonstrated the efficacy of DFOS by applying it to develop a new feed for the high-demand leopard coral grouper (*Plectropomus leopardus*) in Southeast Asia. This system provides an efficient data-driven framework for feed optimization across aquaculture and livestock, contributing significantly to more sustainable and productive farming practices.

## Main

The conservation and restoration of marine and terrestrial environments are important issues for modern society, as outlined in the Sustainable Development Goals (SDGs)^1,2^. ISpecifically, developing and disseminating environmentally friendly aquaculture and livestock farming technologies are urgently required to promote the sustainable use of environmental resources while balancing ecosystem preservation and the needs of an increasing population^3–5^. In this context, the development of feed with low environmental impact is crucial in the advancement of aquaculture and livestock farming technologies. These initiatives promote the efficient growth of aquaculture and livestock farming targets while minimizing their impact on the ecosystem^6^.

The development of an efficient feed is a time-consuming and expensive process that begins with an understanding of the nutritional needs of the target species, followed by the selection and blending of appropriate ingredients^7–9^. Key nutrients, including proteins, fats, amino acids, vitamins, and minerals, significantly affect the growth of farmed animals, and their proper adjustment is crucial to achieve feed efficiency^10,11^. The feed conversion ratio (FCR) is an index of feed efficiency that indicates the ratio of the amount of food consumed to weight gain. Improvements in optimal feed components are required to efficiently increase growth. The FCR varies considerably depending on the target species. However, identifying the optimal combination of these components requires an examination of a large number of possibilities, which makes the process time consuming and cost inefficient.

In information science, the challenge of selecting the optimal candidate from multiple options is known as a black-box optimization problem^12,13^, for which various methods have been proposed. Among them, Bayesian optimization (BO)^14^ is highly effective and has recently been extensively used in various areas such as biology, chemistry, and material development.^15–17^ In BO, a surrogate model is initially constructed to predict the experimental results (in this study, the growth rate when fed a specific feed) from previously obtained data. Next, promising candidates are recommended from the untested options while accounting for the uncertainty of the surrogate model^12,14^. Actual experiments are then conducted on these recommended candidates. Repeating the process of surrogate model development, recommendations, and experiments enables efficient data-driven discoveries. To date, studies have been conducted on prediction models of feed intake and growth for various animals, such as hens, pigs, and fish, with the aim of developing precision feeding systems.^18,19^ In addition, using BO, the feed composition for pigs has been optimized computationally using a model that predicts growth levels rather than through experimentation.^20–22^ However, to the best of our knowledge, no studies have used BO to optimize feed composition while incorporating the results of actual feeding experiments.

Furthermore, applying BO requires determining the specific components in the feed to be optimized, which is a crucial but challenging step. Identifying the components that play critical roles in promoting growth is particularly difficult. Recent advances in omics analyses, such as metabolomic and transcriptomic analyses, have created new opportunities for feed development^23,24^. For example, these techniques can be used to comprehensively analyze the physiological responses of fish and identify key nutritional components essential for growth.^25,26^ However, most studies have focused on evaluating the effects of specific feeds, with limited attention given to leveraging these findings for data-driven development.

In this study, we proposed a data-driven feed optimization system (DFOS) that optimizes feed composition using a data-driven approach by integrating BO and omics analysis. The process of optimizing feed was tested on the aquaculture target species, the leopard coral grouper (*Plectropomus leopardus*)^27^. The DFOS employs a two-stage feed optimization strategy. In the first stage, the proportions of lipids and proteins (the primary components of the feed) were optimized using BO. During this optimization, omics analyses were performed on fish muscle and fecal samples to identify the factors affecting growth by developing a machine learning-based fish growth model. The critical factors identified in the first stage were optimized using BO in the second stage. Furthermore, omics analyses were performed on fish muscle and fecal samples to analyze the factors important for growth, as well as the mechanisms by which feed ingredients affect growth. In this study, we describe the results of applying DFOS to feed development for leopard coral groupers, a grouper species inhabiting East and Southeast Asia that has recently gained attention as a commercially important species in the fisheries industry^28,29^.

## Result

### Data-driven feed optimization system (DFOS)

Figure 1(a) shows an overview of DFOS. DFOS considers a feeding trial to be an input-output system. The ingredients in a feed and their proportions serve as inputs and growth-related indicators such as dairy growth rate and feed efficiency or data from metabolomic and transcriptomic analyses of fish muscle and fecal samples are considered outputs. Subsequently, the feed formulation can be efficiently optimized using BO by analyzing the relationship between the inputs and outputs. DFOS also facilitates the analysis of factors and metabolic pathways necessary for growth.

**Fig. 1.**
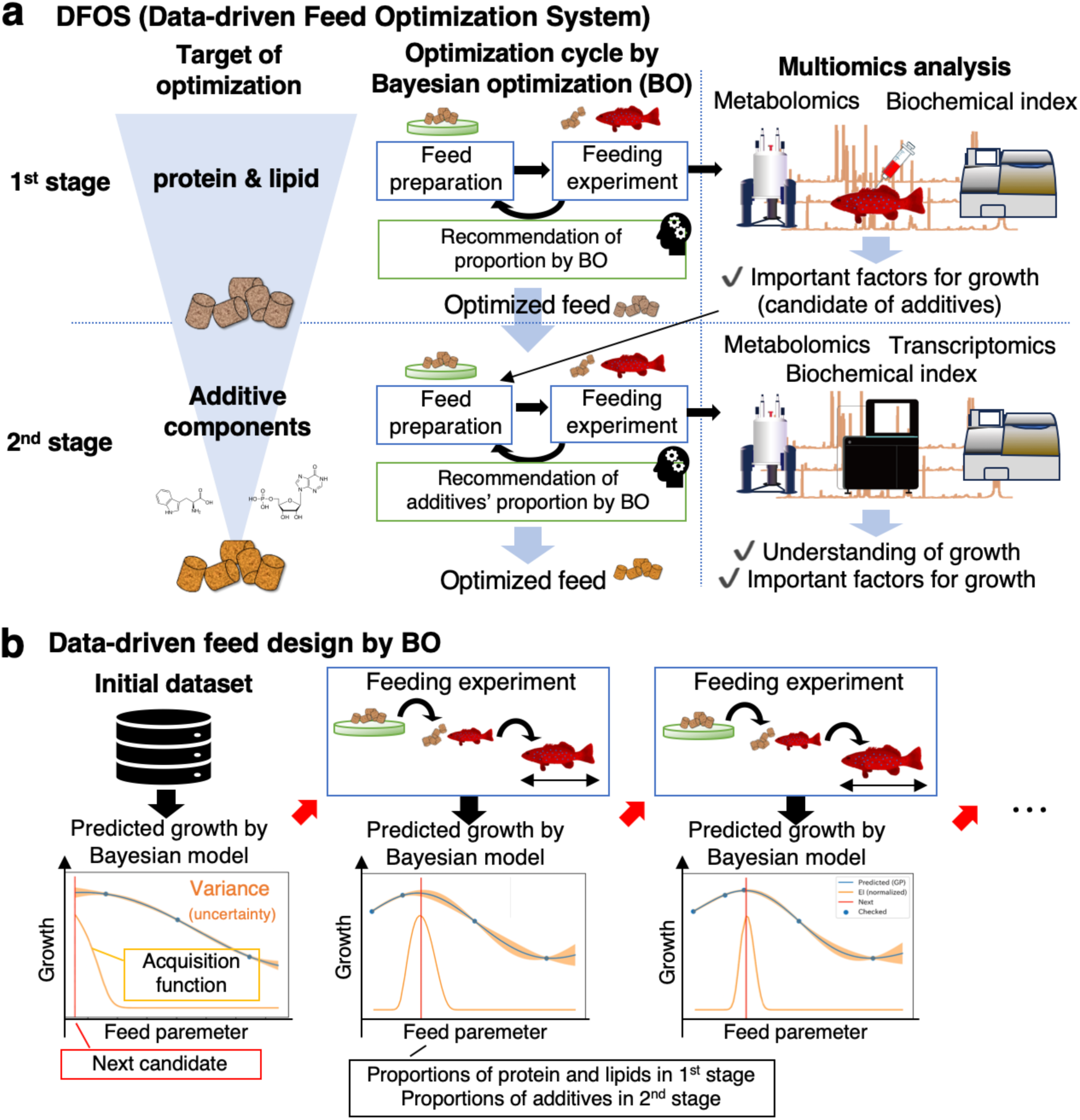
Overview of data-driven feed development with DFOS. (a) In the first stage, DFOS automatically optimizes the proportions of protein and lipid, the primary feed ingredients, using BO. Additionally, from the data obtained in the feed experiments, specific factors potentially related to growth are selected by omics analysis and machine learning. Then, in the second stage, the appropriate combination and proportion of the detailed factors, such as amino acids, obtained in the analysis of the first stage is optimized using BO. (b) Overview of optimization of feed parameters using BO. First, preliminary feeding trials are conducted to obtain feed efficiencies for several diets (initial training set). In BO, a Bayesian model is constructed using Gaussian process regression or other models to predict the feed efficiency and variance for a new feed candidate. Then, an acquisition function is used to select the next feed candidate to be tested. Several candidate feeds may be recommended simultaneously. Feeding experiments are conducted based on the recommended feeds. By repeating this process, the feed parameters are iteratively optimized.

One challenge in data-driven feed optimization is the numerous possible ingredient combinations and proportions in the feed. The DFOS employs a two-stage optimization strategy. First, DFOS optimizes the proportions of key nutrients, protein and lipid, using BO. Specific ingredients that could be added to the feed, such as amino acids, were not adjusted. During this optimization, metabolomic analyses were performed on fish muscle and fecal samples to identify important factors for growth, such as amino acids and compounds. Second, BO is employed to efficiently search for an appropriate combination and proportion of the essential factors selected in the first stage. Figure 1b shows an example of feed efficiency optimization using BO. Gaussian process regression or other methods are used to predict the feed efficiency of a feed that has not yet been tested while estimating its variance. A large variance indicates that the performance of similar feeds has not yet been examined, suggesting significant potential for improvement. An efficient search is achieved by considering the predicted values and variances using an appropriate acquisition function. In addition, omics analysis of feeding trials in the second stage reveals important factors and metabolic pathways involved in growth.

In the following section, we describe the results of feed development for leopard coral groupers using DFOS. With improvements in breeding methods, full-lifecycle aquaculture of the leopard coral grouper was achieved by the Japan Fisheries Research and education Agency (FRA) in 2016.^30^ However, a more efficient feed is needed because the blended feed for sea bream used in aquaculture takes 2.5–3 years for the leopard coral grouper to reach market size.^29^ We aimed to develop a feed with a smaller environmental impact and higher efficiency in which the optimal growth of the leopard coral grouper is achieved with less feed.

### Initial feeding experiments for BO (S1-1 and S1-2)

The protein and lipid content optimization, which are the primary components of the diet, was performed in the first stage. Considering the current advancements in feed development^31,32^ and the practical limitations of feed creation, the most appropriate combination was explored within 40–55% protein and 10–20% lipid content. The initial data must be obtained to enable data-driven optimization through BO. Therefore, six different diets combining three protein levels (45%, 50%, and 55%) and two lipid levels (12% and 16%) were prepared and tested in feeding trials. Details of feed preparation and feeding experiments are provided in the Methods and Supplemental Table S1. The prepared feeds were used in two rearing trials—S1-1 and S1-2—for approximately 8 weeks for juvenile fry.

Table 1 lists the results of S1-1 and S1-2. Although the feeding experiment in one of the tanks fed with P50-L16 in the S1-1 trial was discontinued owing to mass mortality, feeding experiments were conducted successfully in almost all tanks, and survival rates ranged from 94.8% to 100%. The S1-2 trial was large and showed higher daily growth rates and feed efficiency (Table 1) owing to the different periods of the feeding trials. Trials S1-1 and S1-2 confirmed that the daily growth rate and feed efficiency tended to be relatively higher in the plots with higher proportions of proteins and lipids.

**Table 1.** S1-1 and S1-2 feeding trial results. The means and standard deviations of the results of the experiments in the two tanks are shown, except for P45-L16 in S1-1. Values are mean ± SD.

|  |  | P45-L12 | P50-L12 | P55-L12 | P45-L16 | P50-L16 | P55-L16 |
| --- | --- | --- | --- | --- | --- | --- | --- |
| S1-1 | Dairy Growth rates (%/day) | 0.89±0.19 | 0.95±0.08 | 0.86±0.12 | 0.88 | 1.01±0.22 | 0.90±0.10 |
| | Feed efficiency | 0.83±0.21 | 0.84±0.05 | 0.79±0.07 | 0.80 | 0.93±0.23 | 0.82±0.15 |
| | Ave. of initial weights (g) | 27.7±0.99 | 28.0±1.27 | 29.6±2.40 | 29.2 | 27.7±0.99 | 29.2±0.49 |
| | Ave. of final weights (g) | 47.1±6.79 | 49.1±0.14 | 49.2±0.57 | 49.2 | 50.5±4.67 | 49.5±3.82 |
| S1-2 | Dairy Growth rates (%) | 0.13±0.08 | 0.26±0.16 | 0.29±0.13 | 0.28±0.11 | 0.27±0.15 | 0.34±0.10 |
| | Feed efficiency | 0.30±0.15 | 0.51±0.28 | 0.65±0.28 | 0.52±0.24 | 0.57±0.33 | 0.83±0.17 |
| | Ave. of initial weights (g) | 85.9±0.63 | 83.9±5.10 | 86.0±1.92 | 85.9±2.68 | 81.4±1.80 | 84.2±6.65 |
| | Ave. of final weights (g) | 92.4±4.23 | 97.1±2.80 | 102±9.67 | 101±3.42 | 95.2±6.28 | 102±2.52 |

### Protein and lipid optimization by BO (S1-3 and S1-4)

BO recommended promising combinations of feed parameters using data from the initial experiments, and feeding trials were conducted with these recommendations. The feed parameters considered for the recommendations were lipid (10–20%) and protein (40–55%) in 1% increments. Because S1-1 and S1-2 were conducted at different periods, BO recommended candidate formulations that were expected to improve the daily growth rate and feed efficiency based on the data from S1-1 and S1-2, respectively. Candidate formulations are listed in Supplementary Table S2. Based on the results of S1-1 and S1-2, in which higher protein and lipid levels were associated with an improved daily growth rate and feed efficiency, BO recommended higher proportions of both. Two test feeding trials (S1-3 and S1-4) were conducted using the three formulations recommended by BO, as indicated by the asterisks in Supplemental Table S2, and the three formulations, P50-L12, P50-L16, and P55-L16, used in S1-1 and S1-2 as controls. Details of the feeding experiments and feed preparation are provided in the Methods section and Supplemental Table S3.

Table 2 shows the results of the S1-3 and S1-4 feeding trials, including the diets recommended by BO. Mortality during the test period ranged from zero to two fish per tank, and survival rates ranged from 99% to 100% in all tanks. Therefore, diets with a higher percentage of lipids, including P50-L20 and P55-L20, exhibited a higher daily growth rate and feed efficiency, whereas diets with lower lipid content, such as P50-L12, had lower growth performance. However, it was challenging to increase the lipid content using the current preparation method, and the actual percentage in the prepared diets was below the planned value (Supplemental Table S3), empirically indicating that the maximum lipid content should be approximately 17–18%. Additionally, considering the overall results of the feeding trials regarding the protein-lipid ratio, as shown in Supplemental Fig. S1, a ratio of approximately 3 yielded good performance. Based on this ratio, we determined that the P52-L18 ratio was promising. In the subsequent optimization of the second stage, the feed using this ratio, referred to as PL-best, was the base diet, except for S2-1 and S2-2.

**Table 2.**
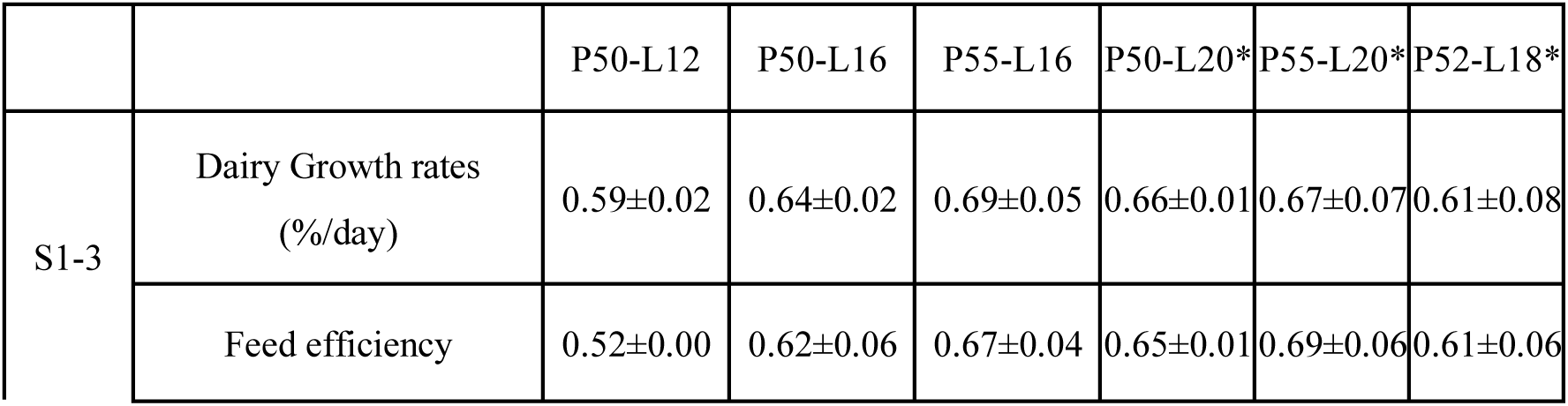
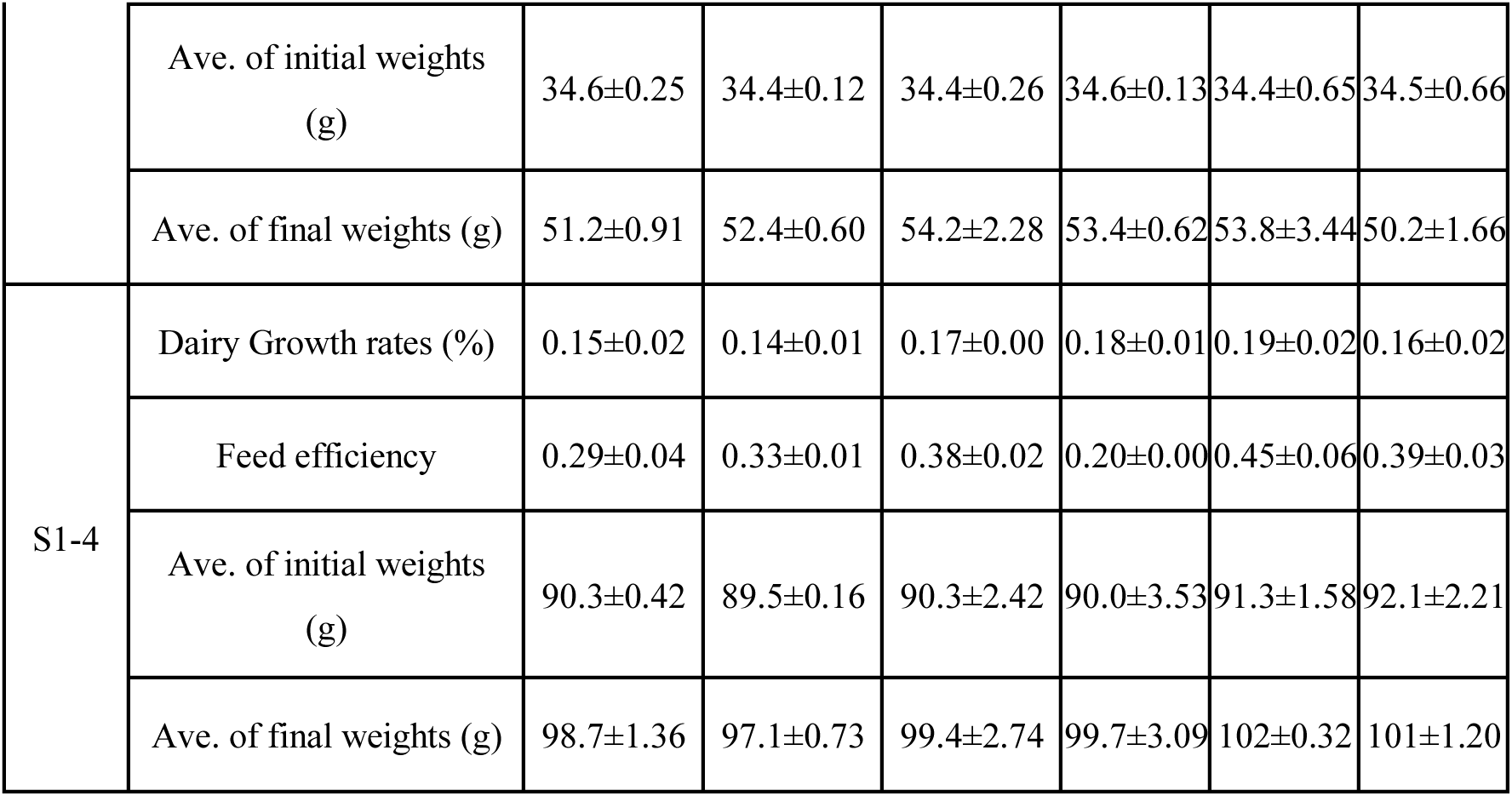
S1-3 and S1-4 feeding trial results. Formulations marked with an asterisk are those recommended by the BO. Other formulations are those used in S1-1 and S1-2 as controls. Values are mean ± SD.

### Analysis of critical factors for growth based on omics analysis

A biochemical analysis using the plasma was conducted to confirm that fish used in the first stage grew healthily without any issues. Fifteen representative indicators of hepatic and urinary functions and carbohydrate, lipid, and protein metabolism were measured (Fig. S2). No significant differences were observed. Although some variation existed, there appeared to be no problematic items.

To identify factors potentially contributing to the growth required in the second stage, metabolome analysis and importance analysis based on the ensemble deep neural network (EDNN) method^24^ were conducted. NMR measurements (two-dimensional J-resolved spectroscopy and 2D-*J*res method) were performed on the fish muscle tissue and feces in experiments S1-1 and S1-2. The details of the analysis are described in the Methods section. We compared the signal intensities in the feed and feces to identify the components that may contribute to growth (Fig. 2(a)). The components shown on the right side of the figure, which were more abundant in the diet but less abundant in the feces, were considered important because of their high consumption by leopard coral groupers. These components included inosine monophosphate (IMP), creatine, lactic acid, sarcosine, and choline. Furthermore, as an alternative analysis approach, a model for estimating the weight using the NMR spectra as the input was constructed using the EDNN method (Fig. 2(b)). The details of the prediction model and analysis method are presented in the Methods section. The developed model was highly accurate, with R^2^=0.77. Using this model, the factors important for prediction were analyzed, as shown in Fig. 2(c), and IMP, arginine, creatine, cystathionine, cysteine, choline, lactate, and others were selected. The relationships between the weight and signal intensities of arginine, choline, cystathionine, creatinine, and lactate are shown in Fig. 2(d).

**Fig. 2.**
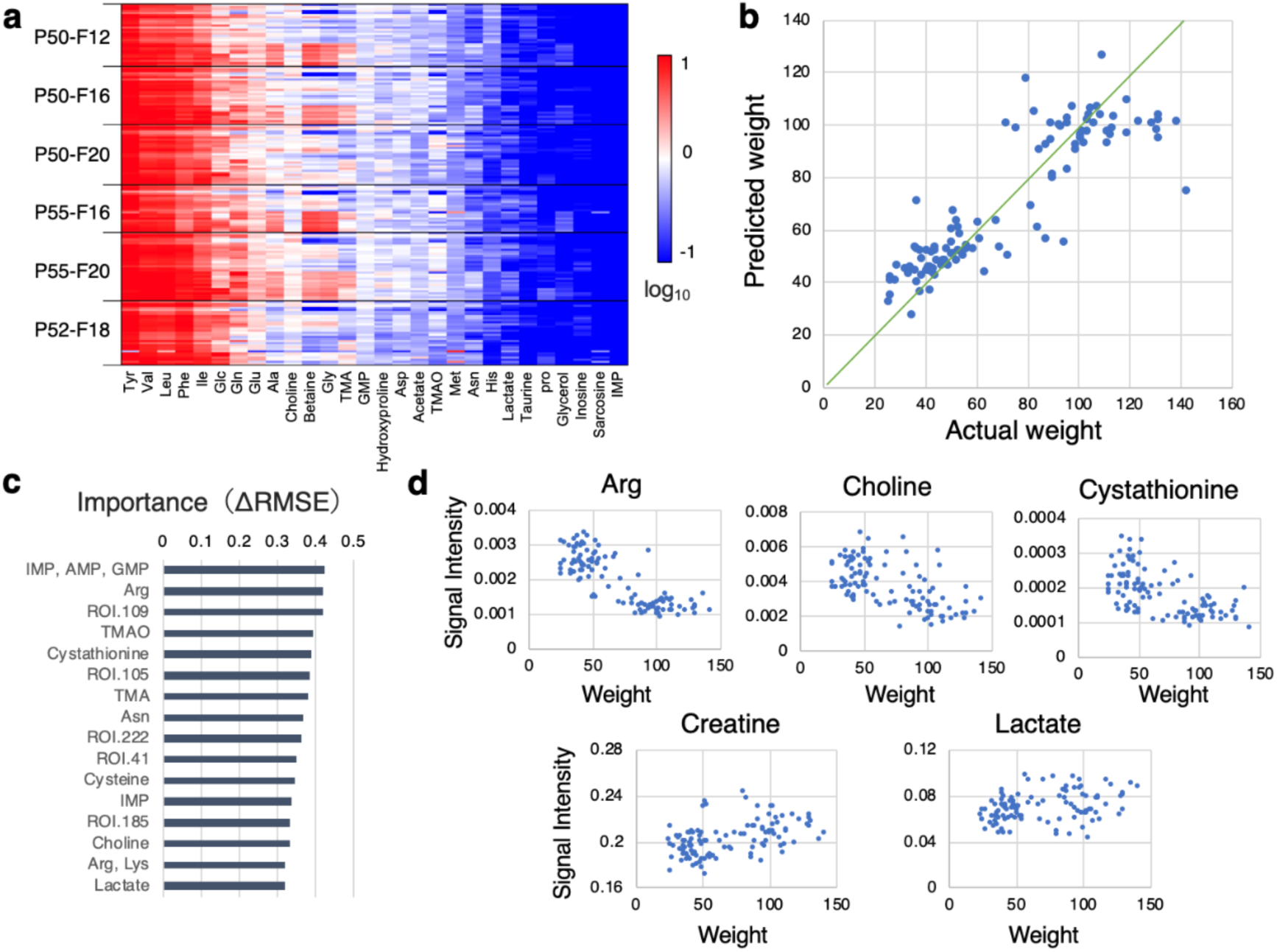
NMR-based metabolome analysis for estimating factors that potentially contribute to growth. (a) Comparison of signal intensities in feed and feces. The NMR signal intensities of feces were divided by the NMR signal intensities of feeds for each peak. The rows are arranged by individual fish, and the columns are arranged by assigned peaks and sorted by average intensity. Colors are scaled from 1.0 to −1.0 using values that were log10 transformed. IMP, lactate, sarcosine, and choline were selected as the most consumed components. (b) Prediction of weight using signals in fish muscle by the EDNN method. (c) Importance values obtained from the EDNN model constructed in (b). IMP, arginine, cystathionine, creatine, and lactate were selected as essential components. (d) Examples of the relationship between weight and factors potentially related to growth.

The data-driven analyses identified various components as potential contributors to growth. Creatine, IMP, lactate, and choline levels were consistently measured. These components were treated as candidate components for diet in the second stage.

### Feeding experiments for optimization of additive components (S2-1 to S2-4)

In the second stage, further optimization was performed by adding creatine, lactate acid, IMP, choline, arginine, and cysteine, which are potentially crucial for growth to the PL-best, protein-, and lipid-optimized diets described above. In addition, GAA, which is a precursor of creatine and glycine, whose importance has been highlighted in previous studies^33–36^ and tryptophan (Trp), which regulates multifunctional effects such as feed intake and growth^37^, were considered candidate ingredients.

Feeding trials S3-1, S3-2, S3-3, and S3-4 were conducted as the initial experiments required for BO to adjust the additives. The detailed results of S3-1 were analyzed in a previous study^38^. In S3-1 and S3-2, designed feeds based on F50-L12 and additives were used, as PL-best could not be prepared in time. In S3-3 and S3-4, the feeds were prepared based on PL-best with additives. Detailed formulations of the diets in S3-1, S3-2, S3-3, and S3-4 are described in Supplemental Tables S4, S5, S6, and S7, respectively. Feeding experiments for each diet were conducted in three small tanks at a constant water temperature to ensure stable results in subsequent trials. The details of the feeding experiments are provided in the Methods section.

The results of experiments S3-1, S3-2, S3-3, and S3-4 are presented in Table 3. Some diets containing IMP, glycine, choline, and tryptophan outperformed the control (P50-L12 and PL-best) and were superior in terms of daily growth rate and feed efficiency. However, diets containing cysteine, arginine, GAA, and creatine showed similar or lower performance than the control. Therefore, in subsequent stages, we optimized the combination and ratio of IMP, glycine, choline, and tryptophan.

**Table 3.**
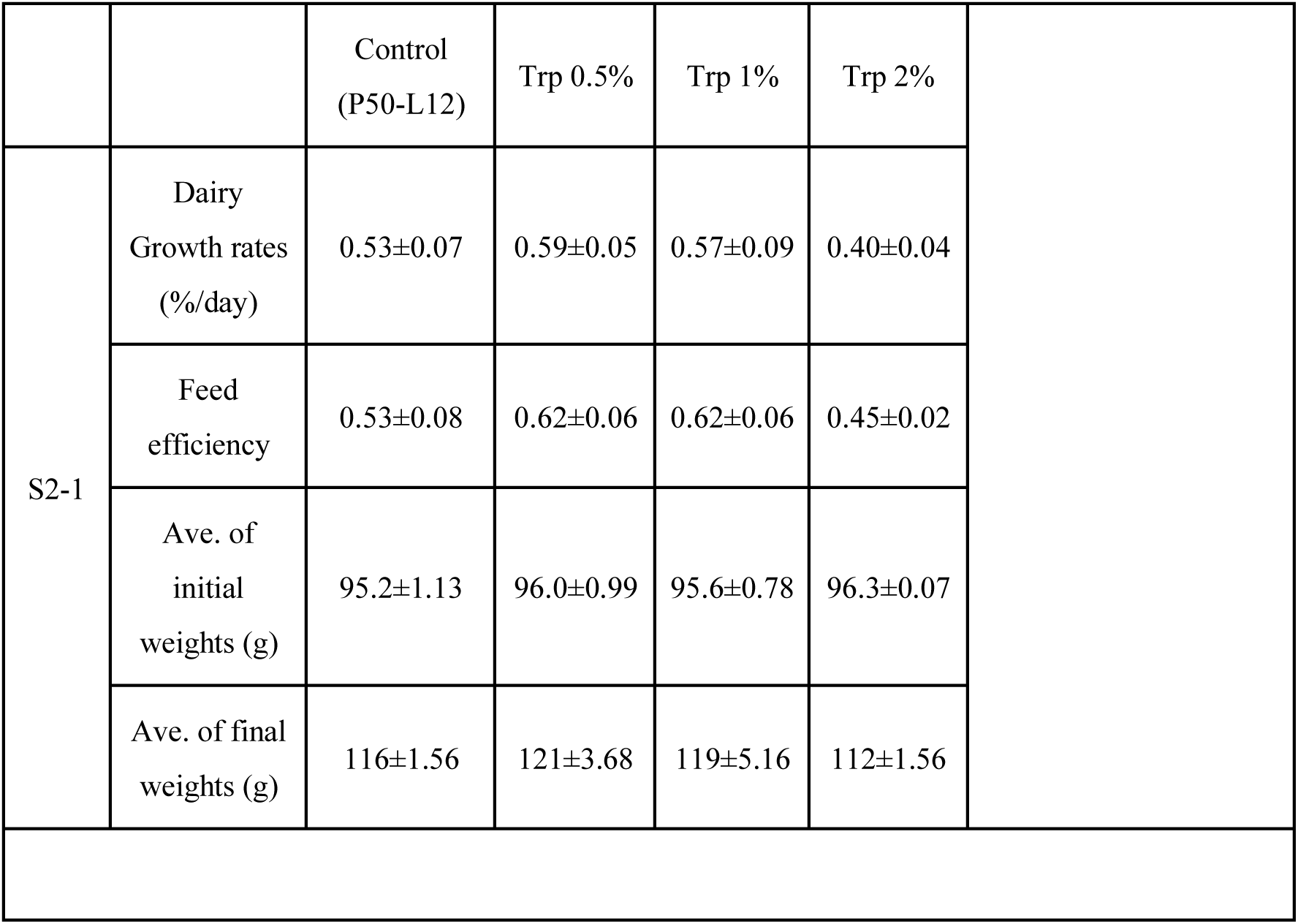
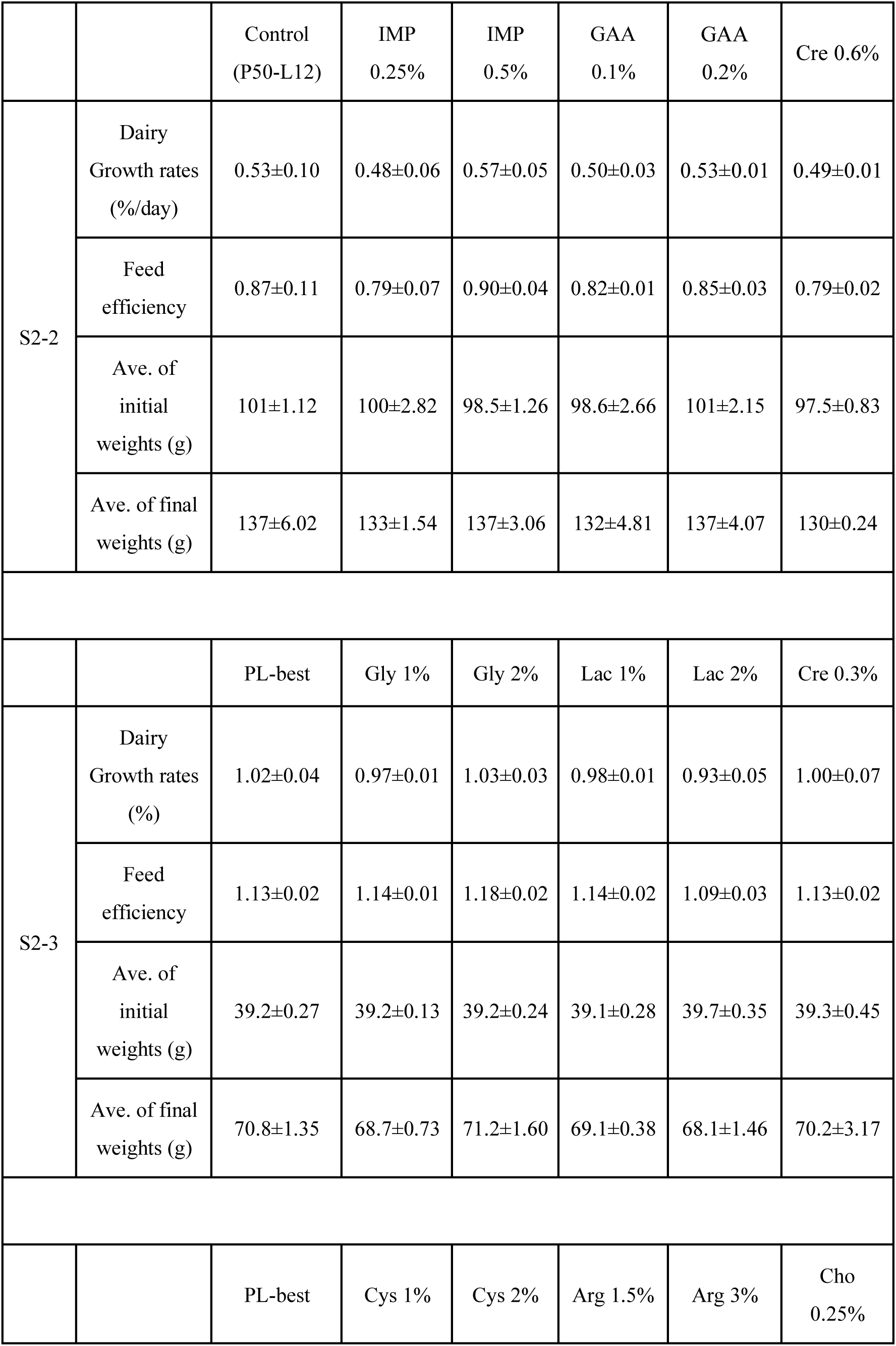
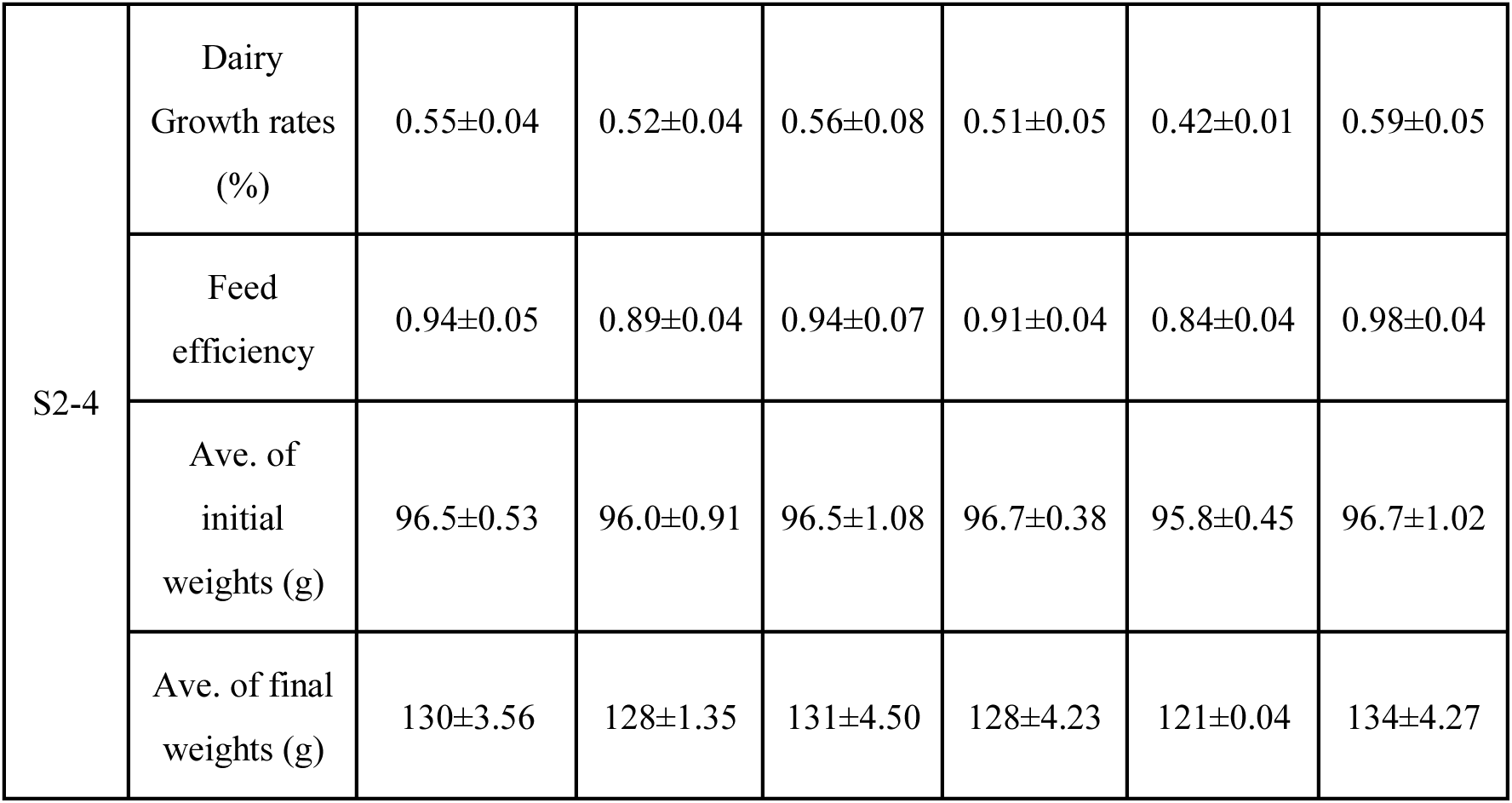
Formulation of S2-1, S2-3, S2-3, and S2-4 feed trials as initial experiments for additive optimization and their results of feeding experiments. Values are mean ± SD.

### Optimization of additives by BO (S2-5 and S2-6)

Based on the results of these feeding trials, two BO cycles were conducted to optimize the additives. The details of the BO during the additive optimization are described in Section 4.8. The candidate additives and their ranges were tryptophan (0%, 0.25% and 0.5%), IMP (0%, 0.25%, 0.5%, 0.75%, and 1%), glycine (0%,1%, 2%, 3%, and 4%), and choline (0%, 0.25%, 0.5%, 0.75%, and 1%), resulting in 375 combinations. In the first recommendation of additives for BO, feed components BO1, BO2, and BO3, shown in Supplemental Table S8, were suggested as candidates to maximize feed efficiency, daily growth rate, and the product of feed efficiency and daily growth rate. The feed trial S2-5 was conducted using PL-best as a control, choline 0.5% (which could not be tested until S2-4), and BO-recommended BO1, BO2, and BO3. Details of feed formulations are presented in Supplemental Table S9. Table 4 lists the results for S2-5. The feeding experiments showed that BO1 and BO2 demonstrated a slight improvement in feed efficiency, although no significant differences were observed in the daily growth rate.

**Table 4.** S2-5 feeding trial results. Values are mean ± SD.

|  |  |  |  |  |  |  |
| --- | --- | --- | --- | --- | --- | --- |
|  |  | PL-best | Cho-0.5 | BO1 | BO2 | BO3 |
| S2-5 | Dairy Growth rates (%/day) | 0.47±0.06 | 0.40±0.05 | 0.46±0.05 | 0.46±0.01 | 0.44±0.05 |
|  | Feed efficiency | 0.84±0.04 | 0.78±0.06 | 0.85±0.03 | 0.86±0.03 | 0.81±0.07 |
|  | Ave. of initial weights (g) | 138±1.97 | 136±1.84 | 138±4.30 | 141±4.69 | 139±2.41 |
|  | Ave. of final weights (g) | 180±6.90 | 170±2.18 | 178±4.24 | 182±7.52 | 178±7.38 |

The second cycle of additive optimization using BO was conducted as trial S2-6 based on the 2-5 feeding trial results for the additives. Formulations BO4, BO5, BO6, and BO7 recommended by the BO are shown in Supplemental Table S10. Table 5 presents the S2-6 feeding trial results. PL-best was used as a control. BO7 demonstrated a higher daily growth rate and feed efficiency than the control.

**Table 5.** S2-6 feeding trial results. Values are mean ± SD.

|  |  |  |  |  |  |  |
| --- | --- | --- | --- | --- | --- | --- |
|  |  | PL-best | BO4<br>(Multi1) | BO5<br>(Multi2) | BO6<br>(Multi3) | BO7<br>(FE 3) |
| S2-6 | Dairy Growth rates (%/day) | 1.02±0.02 | 1.02±0.09 | 0.97±0.04 | 1.02±0.05 | 1.07±0.05 |
|  | Feed efficiency | 1.15±0.01 | 1.13±0.05 | 1.09±0.01 | 1.10±0.05 | 1.17±0.02 |
|  | Ave. of initial weights (g) | 42.1±0.33 | 42.2±0.42 | 41.7±0.15 | 42.2±0.40 | 41.8±0.35 |
|  | Ave. of final weights (g) | 74.6±1.43 | 74.9±4.10 | 71.8±1.66 | 74.9±2.64 | 76.0±1.37 |

### Omics analysis in the second stage

Biochemical indices, metabolomics, and transcriptomic analyses were conducted to understand the growth of the fish used in the S2-5 and S2-6 trials and fed the developed diets. The biochemical index was measured using plasma; the results are shown in Fig. S2. Some values exhibited different trends for S2-5 and S2-6. However, no significant differences were observed between controls. Regarding lipid metabolism, the levels of TCHO-PIII and TG-PIII showed a decreasing trend in the control and BO7 groups. Additionally, metabolomic analysis of the results in S2-5 and S2-6 was conducted in the same manner as in Stage 1. Figure 3 shows a comparison of the signal intensities in the feed and fecal samples. Although many different metabolites existed in the distribution during stages 1 and 2, the difference between S2-5 and S2-6 appeared to be relatively small. The effects of additives such as Gly and Trp were evidently observed.

**Fig. 3.**
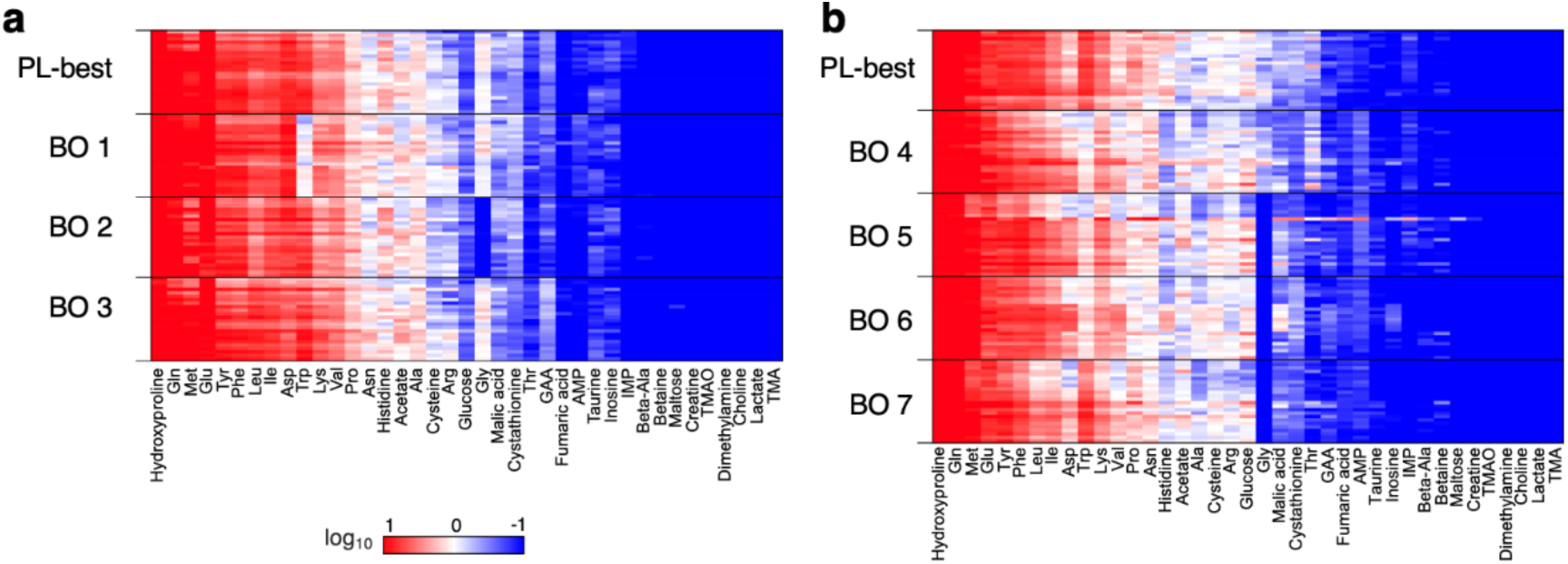
Results of metabolomic analysis S2-5 (a) and S2-6 (b). The NMR signal intensities of feces were divided by the NMR signal intensities of feeds for each peak. The rows are arranged by individual fish, and the columns are arranged by assigned peaks and sorted by average intensity. Colors are scaled from 1.0 to −1.0 using values that were log10 transformed.

Furthermore, a comprehensive mRNA expression analysis was performed to determine the efficiency of the new diets. The cDNA libraries constructed from the hepatopancreas of the leopard coral grouper were sequenced using Illumina NextSeq 500. Sequence data were deposited in the DDBJ Sequence Read Archive under accession nos. PRJDB19825. Each sample yielded an average of approximately 12.2 million sequence reads. Almost all clean reads (99.9% of raw reads) remained and were mapped to the reference sequences at an average rate of 90.0%. Discrimination analysis of the PCA was performed on all groups in the S2-6 trial (Fig. S2). Samples in the same group were widely scattered along components 1–3 with no apparent separation between the groups. BO7, which exhibited a positive growth trend during the rearing experiment, was compared to the control group (PL-best) in the following analysis. The differential expression analysis results are presented in Table 6. Compared to the control group, 123 differentially expressed genes (DEGs) were identified in BO7. Subsequently, pathway analysis was performed, and Table 6 presents the results that utilized the fold change value ˃ 1.5. The number of genes involved is expressed as a score^39^. The SWI/SNF and IGF receptor pathways exhibited high scores in the BO7 group. The keywords cell-cell signaling, cell population proliferation, and cell growth were extracted in the Bio-Event analysis.

**Table 6.** Pathway and enrichment analyses results using transcriptome analysis (S2-6 Ctrl vs BO7). The score was calculated from probabilities based on the hypergeometric distribution.

| Pathway analysis |  |  |
| --- | --- | --- |
| rank | name | score |
| 1 | SWI/SNF signaling pathway | 26.346 |
| 2 | IGF receptor signaling pathway | 10.687 |
| 3 | Transcriptional regulation by BLIMP-1 | 9.573 |
| 4 | Transcriptional regulation by FOXO | 9.078 |
| 5 | Lipid-transfer protein signaling pathway | 8.698 |
| 6 | Transcriptional regulation by PPARγ | 8.545 |
| 7 | Insulin receptor signaling pathway | 7.632 |
| 8 | Transcriptional regulation by Ikaros | 6.492 |
| 9 | Transcriptional regulation by MRF | 5.156 |
| 10 | Tetrahydrobiopterin metabolism | 5.058 |
| Bio-Event |  |  |
| rank | name | score |
| 1 | cell-cell signaling | 16.362 |
| 2 | regulation of cell population proliferation | 16.292 |
| 3 | regulation of cell growth | 15.892 |
| 4 | cell population proliferation | 15.015 |
| 5 | cell growth | 14.985 |
| 6 | regulation of growth | 13.895 |
| 7 | positive regulation of cell population proliferation | 12.849 |
| 8 | leukocyte activation | 12.643 |
| 9 | growth | 12.424 |
| 10 | regulation of apoptotic process | 12.239 |

## Discussion

DFOS optimized the protein-lipid ratio in the first stage and additives in the diets in the second stage for the leopard coral grouper. Figure 4a and b show the optimization of proteins and lipids using BO in the first stage. For comparison, the results of each trial were scaled such that the value of P50-L16 was set to 1, and the averages of S1-1 and S1-2, as well as S1-3 and S1-4, are shown. Higher lipid and protein content corresponded to better growth, and BO was successfully recommended as a promising feed candidate. In addition, the metabolome analysis (Fig. 3) identified factors that might contribute to growth, facilitating additive adjustment in the second stage. Figure 4c and d show the results of the two BO cycles for additive optimization. These results were scaled with the PL-best set to 1 for comparison. Although no notably improved growth feeds were recommended owing to the already optimized protein and lipid levels, BO suggested diets containing multiple additives, including BO7, which improved the daily growth rate and feed efficiency. These results indicate the effectiveness of DFOS-data-driven feed optimization.

**Fig. 4.**
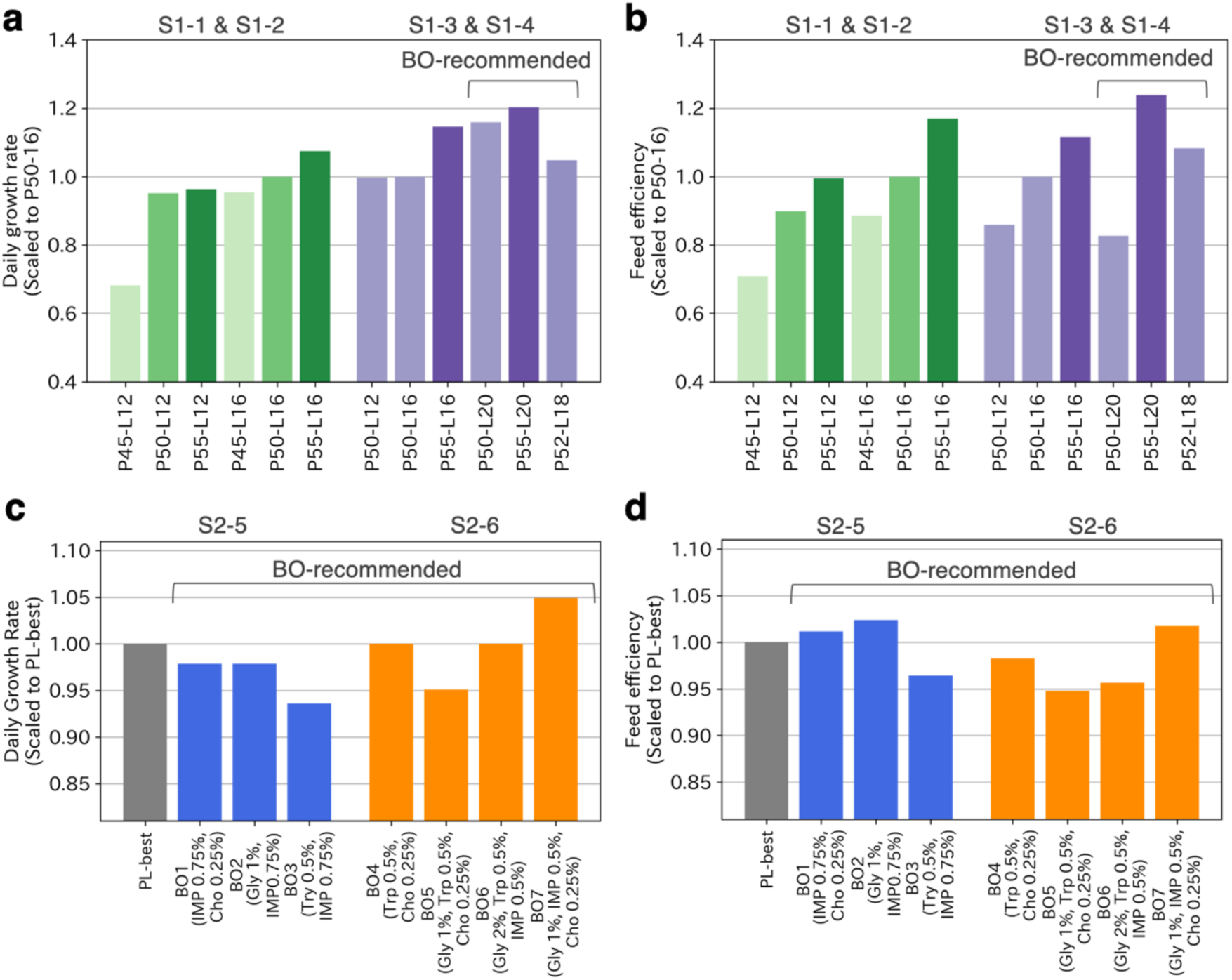
Summary of results of feed recommendation by BO. (a) and (b) Daily growth rate and feed efficiency optimized with BO for the proportion of protein and lipid in the first stage. Darker colors indicate a higher percentage of protein. (c) and (d) Results in the second stage, where the combination of additives and their proportions were optimized by BO.

The transcriptomic analysis results suggested why BO7 improved. Multivariate analysis using PCA did not reveal distinct differences among the groups; however, pathways and their related analyses yielded two keywords. The IGF receptor signaling pathway, transcriptional regulation by FOXO, and the insulin receptor signaling pathway are important in growth factors and cell proliferation^40–45^. Therefore, BO7 is believed to exhibit the optimal growth rate.

Several studies have been conducted on the feed for leopard coral groupers. The selection of nutrient-enhanced rotifers and feeding environments, such as water temperature and lighting conditions, have been tested^27,46–48^. For juveniles, the optimal feed protein levels were examined ^31,32^, and the effect of astaxanthin additive was verified^49^. The basic physiological features have been reported using comprehensive analysis methods^25,26^. Focusing on the amino acid levels in the feed is also important. The addition of amino acids is expected to have desirable effects. Recently, optimal concentrations of specific nutrients in feed, such as tryptophan, lysine, and vitamins, have been identified^38,50,51^. However, studies on the optimization of comprehensive nutrition using BO are lacking. In addition, although a study exists on optimizing pig feed composition using a computational model^20^, this study is the first to conduct a feeding experiment to demonstrate the effectiveness of feed optimization using BO. In addition, although most feed development projects tend to optimize a single component, optimizing multiple parameters such as proteins, lipids, and additives simultaneously using BO is possible. DFOS is expected to reduce the time and cost of feed development.

From an optimization method perspective, DFOS has some room for improvement. As shown in the Supplemental Tables of the feed formulations, some discrepancies were observed between the planned formulation and the actual produced diets. This may negatively affect the BO’s optimization, and minimizing these discrepancies is necessary. Notable fluctuations were observed between tanks in the feeding experiments, possibly owing to the sensitivity and vigilance of leopard coral groupers, which make them easily affected by their environment. More stable rearing experiments are expected to enable a more effective search for feed. Furthermore, in this study, the experimental conditions, such as the number of tanks and water temperature control, were changed between stages 1 and 2 to stabilize the feeding trial results. Although conducting experiments under consistent conditions is preferable, developing a BO-based method that considers variations in feeding conditions is also an issue for future studies.

## Conclusion

We developed DFOS, a data-driven feed optimization system. The proposed system optimizes diet components and additives based on the evidence from feeding experiments. By integrating omics analysis, we presented a novel strategy for data-driven optimization of fine feed ingredients. In addition, by using the BO, the system enables automated searches. To demonstrate the effectiveness of DFOS, we applied the proposed system to develop diets for the leopard coral grouper and successfully optimized the diets with improved feed efficiency and daily growth rates. Furthermore, omics analysis provided insights into essential growth mechanisms. DFOS, as a general data-driven framework, can be applied to other species and contributes to the development of environmentally friendly and cost-effective feeds.

## Methods

### Development of feed

In the present study, various diets with different compositions were prepared. Details of each diet formulation are provided in the Supporting Information: S1-1 and S1-2 in Tables S1, S1-3, S1-4 in Table S3, S2-1 in Table S4, S2-2 in Table S5, S2-3 in Table S6, S2-4 in Table S7, S2-5 in Table S9, and S2-6 in Table S11. For compositional analysis, the diets were heated at 105 °C for 15 h, and moisture content was determined by measuring the weight loss. The nitrogen content was evaluated using the semi-micro Kjeldahl method, and crude protein was calculated by multiplying the nitrogen content by 6.25 (N × 6.25). Crude lipid content was determined using the Folch method, and ash content was ashed at 600 °C for 5 h.

### Ethics statement

Experiments were conducted in accordance with the Principles and Guidelines for the Care and Use of Live Fish and the Guidelines for Animal Experimentation of the National Research Institute of Fisheries Science, Fisheries Research Agency. All experimental methods were approved by the Animal Experimental Council (AEC/NRIFS) of the National Research Institute of Fisheries Science, Fisheries Research Agency. Fish were anesthetized with 2-phenoxyethanol (Wako, Osaka, Japan), and efforts were made to minimize suffering.

### Feeding experiment and growth parameters

All feeding experiments were conducted at Yaeyama Laboratory, Fisheries Technology Institute, Japan Fisheries Research and Education Agency. Juvenile leopard coral groupers were used in all experiments, but their age in months varied. The volume of the tanks and the stocking density were set according to the growth conditions of the fish. The volume of the first stage and S2-1 tanks was 2.4 kL (1 kL of seawater), and the stocking density was around 10 kg/kL. Two tanks were used for each feed treatment. In the second stage, excluding S2-1, the volume of the tanks was 0.2 kL (0.15 kL of seawater inside), and the fish were housed in three tanks for each feed. The stocking density was around 10 kg/kL for S2-2, −4, and −5 and 5 kg/kL for S2-3 and −6. Detailed fish sizes are provided in the tables for each experimental result. The UV-sterilized and sand filtered seawater was aerated. The total length and weight of the fry were measured before and after the feeding experiments. Every seven days in morning, fish fecal were collected by the syphon. Collected fish fecal samples were stocked in the deep freezer till analyzing. Although the feeding experiments were conducted under the same conditions as possible, some conditions were not identical owing to time, space, or budget constraints. In the first stage, the water temperature was ambient. In the second stage, the water temperature was maintained at 25°C. Growth parameters, dairy growth rate and feed efficiency were calculated as follows:

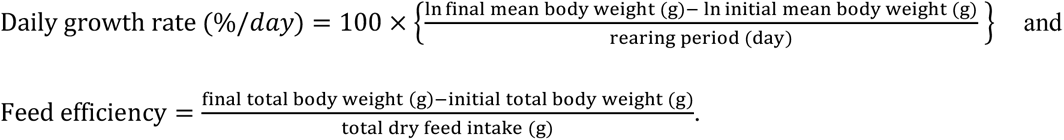

### Metabolome analysis

Low molecular metabolites were measured using an AVANCE II 700 MHz NMR spectrometer (Bruker BioSpin, Rheinstetten, Germany) with a 2D *J*-resolved (2D*J*-res) pulse program. Lyophilized muscle, feed, and feces samples were shaken (1,400 rpm) with potassium phosphate (KPi) buffer solution in D2O, containing 1 mmol/L sodium 2,2-dimethyl-2-silapentane-5-sulfonate (DSS) for 15 min at 65 °C.

After centrifugation at 14,000 rpm for 5 min, the extracted supernatants were transferred to an NMR tube. Following measurement, the spectra were peak-picked using the rNMR software, which identified 225 peak intensities common to the samples, and 119 were annotated.^52^ SpinAssign^53^ and SpinCouple^54^ were used to annotate the peaks. The intensity of each peak was calculated by aligning the sum of the peak intensities of each spectrum, and was used to compare the intensities of each substance and to train a prediction model.

### Measurement of plasma biochemical index

Plasma was applied to the following slides: alanine aminotransferase (GPT/ALT-PⅢ), alkaline phosphatase (ALP-PⅢ), amylase (AMYL-PⅢ), aspartate aminotransferase (GOT/AST-PⅢ), inorganic phosphorus (IP-P), leucine aminopeptidase (LAP-P), albumin (ALB-PⅢ), creatinine (CRE-PⅢ), glucose (GLU-PⅢ), total cholesterol(TCHO-PⅢ), total protein (TP-PⅢ), total bilirubin (TBIL-PⅢ), and triglyceride(TG-PⅢ), urea-nitrogen (BUN-PⅢ), and uric acid (UA-PⅢ). The slides were loaded to the Fuji DRI-CHEM NX500 (Fujifilm, Tokyo, Japan).

### Transcriptomic analysis

Total RNA was extracted from the hepatopancreas using Direct-Zol (Zymo Research, Irvine, CA, USA) according to the manufacturer’s protocol. RNA quality was evaluated based on the proportion of rRNA using an Agilent 2200 TapeStation system (Agilent Technologies, Palo Alto, CA, USA). Complementary DNA libraries were constructed using the llumina Stranded mRNA Prep, Ligation (Illumina, San Diego, CA, USA). Sequencing was performed using an Illumina Nextseq500 platform equipped with a 75 bp single-end module. Demultiplex and multi-index adaptors were trimmed from the sequenced raw reads using Illumina software (bcl2fastq). Removal of low-quality sequences and following de novo assembly were performed using CLC Genomics Workbench 21.1 (Qiagen, Hilden, Germany). The parameter of low-quality sequence was 0.05, and removal of ambiguous nucleotides was 2. The read mapping analysis was performed using the reference sequence of *P. leopardus*. Clean reads were aligned with the reference genome using the CLC Genomics Workbench, and expression values and transcripts per million (TPM) were calculated. Pathway and Bio-Event analysis was performed by KeyMolnet lite software (KM data, Tokyo, Japan).

### Development of a fish weight prediction model

An ensemble deep neural network (EDNN) was used to predict fish weight using metabolomic data.^24^ This deep neural network-based model could improve accuracy through result integration by ensemble and indicate the importance of variables to the model. Owing to the inability to install ‘mxnet’ for R as used in the study, a similar analysis was performed using Python (3.9.7) and the Sequential model of Keras (2.6.0). Using body weight of fish as the objective variable and NMR peaks as the explanatory variable, cross-validation was performed using 10-fold with samples divided into 10 groups based on number. The importance of variables is shown as ΔRMSE, indicating the influence each variable has on RMSE. The model conditions were as follows: the classifier used for the ensemble was a 5-layer deep neural network, the number of classifiers was 300, the number of variables for each classifier was the square root of the total number of all peaks, and the number of classifiers used to combine the results was set to an RWC of 0.8.

### Bayesian optimization

BO is a methodology that identifies promising candidates with the desired property in the fewest number of experiments possible^13,14^. In the optimization of the first stage, the candidate parameters are combinations of protein and lipid proportions. To make a recommendation with BO, a prediction model needs to be built, so several initial pairs of data on feed formulations and growth rates, in this case daily growth rates and feed efficiency, are required. Prediction models were built to estimate daily growth rate and feed efficiency from S1-1 and S1-2 data based on Gaussian process regression^55^. In this study, model construction and the recommendation of candidates, described below, were performed using PHYSBO^56^, a Bayesian optimization package in which hyperparameters of the model are automatically optimized based on maximizing Type II likelihood^55^. Next, the acquisition function was used to recommend the next candidate to be experimented with. In this study, the expected improvement (EI)^12^, one of the most commonly used acquisition functions, was used.

In the optimization of the second stage, feed candidates were combinations of added amounts of dietary tryptophan, IMP, glycine, and choline. Using the data from S2-1, S2-2, S2-3, and S2-4, Gaussian process regression models were constructed with feed efficiency, daily growth rate, and their product as objective variables, respectively; three promising candidates were selected for each objective variable with EI as the acquisition function. The results for S2-1, S2-2, S2-3, and S2-4 showed large differences because of the different feeding periods and fish sizes used in each test. Thus, all data were unified by scaling the mean feed efficiency and daily growth rate in the control for each test to 1 and 1%, respectively. The expected values and variances shown in Supplemental Table S8 represent the values at which this unification occurred. PHYSBO was used to implement BO, same as in the first stage.

## Acknowledgement

This work was supported and founded by the Fisheries Agency of Japan (Research project for the transformation of the aquaculture industry into a growth industry in 2019−2023). The authors wish to thank Dr. Masahiro Kobayashi for organizing the project Dr. Ryo Kimura and Dr. Masahiko Koiso for supervision and administration, and Ms. Kazuyo Iida for data acquisition support.

## Data availability

The data and codes generated in the present study are available from the corresponding authors upon reasonable request.

