## Supporting Information for "Data-Driven Feed Optimization for Sustainable Aquaculture through Integrated Omics Analysis and Two-stage Bayesian optimization"

Table S1. Formulation and proximate composition of the test diets for S1-1 and S1-2.

| | P45-L12 | P50-L12 | P55-L12 | P45-L16 | P50-L16 | P55-L16 |
|---|---|---|---|---|---|---|
| Ingredients (g/kg) | | | | | | |
| Fish meal (Anchovy/Sardine) | 440 | 510 | 550 | 440 | 510 | 570 |
| Krill meal | 90 | 100 | 110 | 90 | 100 | 110 |
| Wheat flour | 193 | 228 | 273 | 203 | 233 | 218 |
| Tapioca starch | 200 | 90 | 0 | 150 | 50 | 0 |
| Fish oil | 20 | 15 | 10 | 60 | 50 | 45 |
| Vitamin mixture[1] | 27 | 27 | 27 | 27 | 27 | 27 |
| Mineral mixture[2] | 30 | 30 | 30 | 30 | 30 | 30 |
| Proximate composition (% dry matter basis) | | | | | | |
| Moisture | 4.7 | 3.2 | 4.0 | 3.7 | 3.4 | 3.5 |
| Crude protein (%) | 43.0 | 49.1 | 54.1 | 43.2 | 48.4 | 53.9 |
| Crude Lipid (%) | 11.8 | 12.7 | 13.4 | 15.5 | 16.0 | 17.0 |
| Ash | 10.0 | 10.9 | 11.7 | 9.9 | 10.9 | 11.7 |

[1] Vitamin mix (amounts per 100 g of diet): thiamine nitrate, 18.5 mg; riboflavin, 12 mg; pyridoxine HCl, 14.6 mg; nicotinamide, 44.6 mg; Ca patothenate, 32.6 mg; myo-inositol, 120 mg; biotin, 0.3 mg; folic acid, 0.9 mg; menadione bisulfite sodium, 4.6 mg; a-tocopherol acetate, 12 mg; cyanocobalamin, 30 µg; Rovimix stay-C35, 200 mg; choline chloride, 500 mg; VA palmitate, 440 IU; vitamin D. 90 IU.

[2] Mineral mix (amounts per 100 g of diet): $MnSO_4$, 7.8 mg; iron(II) fumarate, 150.0 mg; $CoSO_4$, 0.12 mg; $CuSO_4$, 1.5 mg; $ZnSO_4$, 17.1 mg; KI 0.15 mg; $MgSO_4$, 510.0 mg; aluminum hydroxide, 16.3 mg; carcium lactate, 150.0 mg; $Ca(H_2PO_4)_2$, 2.1 g.

Table S2. The formulations recommended by BO from S1-1 and S1-2 and their predicted values. Objective is the objective variable for recommendation by BO. Items marked with an asterisk were employed in the S1-3 and S1-4 feeding trials.

| Objective | Protein | Lipid | Predicted value (variance) |
| --- | --- | --- | --- |
| Dairy Growth rates (S1-1)* | 50% | 20% | 0.74 (0.22) |
| Feed efficiency (S1-1)* | 55% | 20% | 0.95 (0.0047) |
| Dairy Growth rates (S1-2) | 48% | 20% | 0.14 (0.15) |
| Feed efficiency (S1-2)* | 52% | 18% | 0.45(0.0045) |

Table S3. Formulation and proximate composition of the test diets for S1-3 and S1-4. The detailed compositions of the Vitamin and Mineral mixtures are described in Table S1.

| | P50-L12 | P50-L16 | P55-L16 | P50-L20* | P55-L20* | P52-L18* |
|---|---|---|---|---|---|---|
| Ingredients (g/kg) | | | | | | |
| Fish meal (Anchovy/Sardine) | 530 | 530 | 600 | 530 | 600 | 550 |
| Krill meal | 110 | 110 | 120 | 110 | 120 | 110 |
| Wheat flour | 178 | 173 | 138 | 178 | 143 | 158 |
| Tapioca starch | 110 | 80 | 40 | 40 | 0 | 60 |
| Fish oil | 15 | 50 | 45 | 85 | 80 | 65 |
| Vitamin mixture | 27 | 27 | 27 | 27 | 27 | 27 |
| Mineral mixture | 30 | 30 | 30 | 30 | 30 | 30 |
| Proximate composition (% dry matter basis) | | | | | | |
| Moisture | 9.1 | 4.2 | 5.2 | 5.0 | 5.6 | 5.0 |
| Crude protein (%) | 50.9 | 50.3 | 55.6 | 50.9 | 55.4 | 52.6 |
| Crude Lipid (%) | 10.7 | 14.5 | 15.1 | 18.0 | 18.5 | 16.7 |
| Ash | 11.8 | 11.7 | 12.8 | 11.8 | 12.8 | 12.0 |

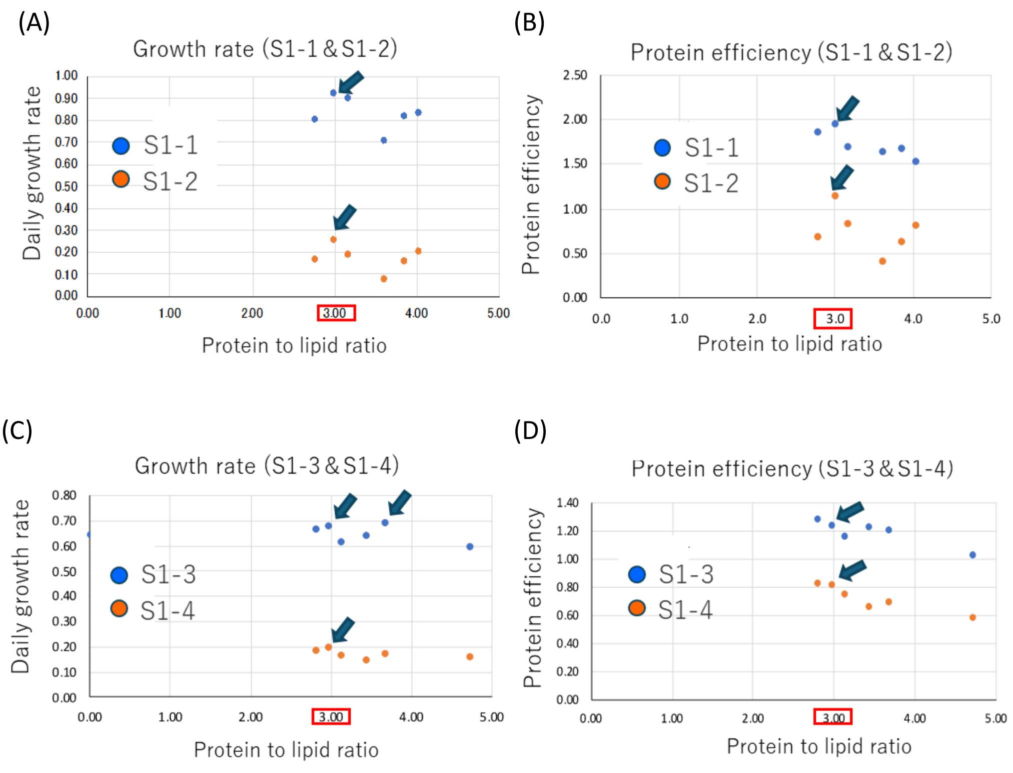

Figure S1. The results of feeding expriments of stage 1. All graphs utilize the protein to lipid ratio as the horizontal axis. Arrowheads indicate the optimal outcome at each stage (A) daily growth rate of stage 1-1 and 1-2, (B) protein efficiency of stage 1-1 and 1-2, (C) growth rate of stage 1-3 and 1-4, (D) protein efficiency of stage 1-3 and 1-4.

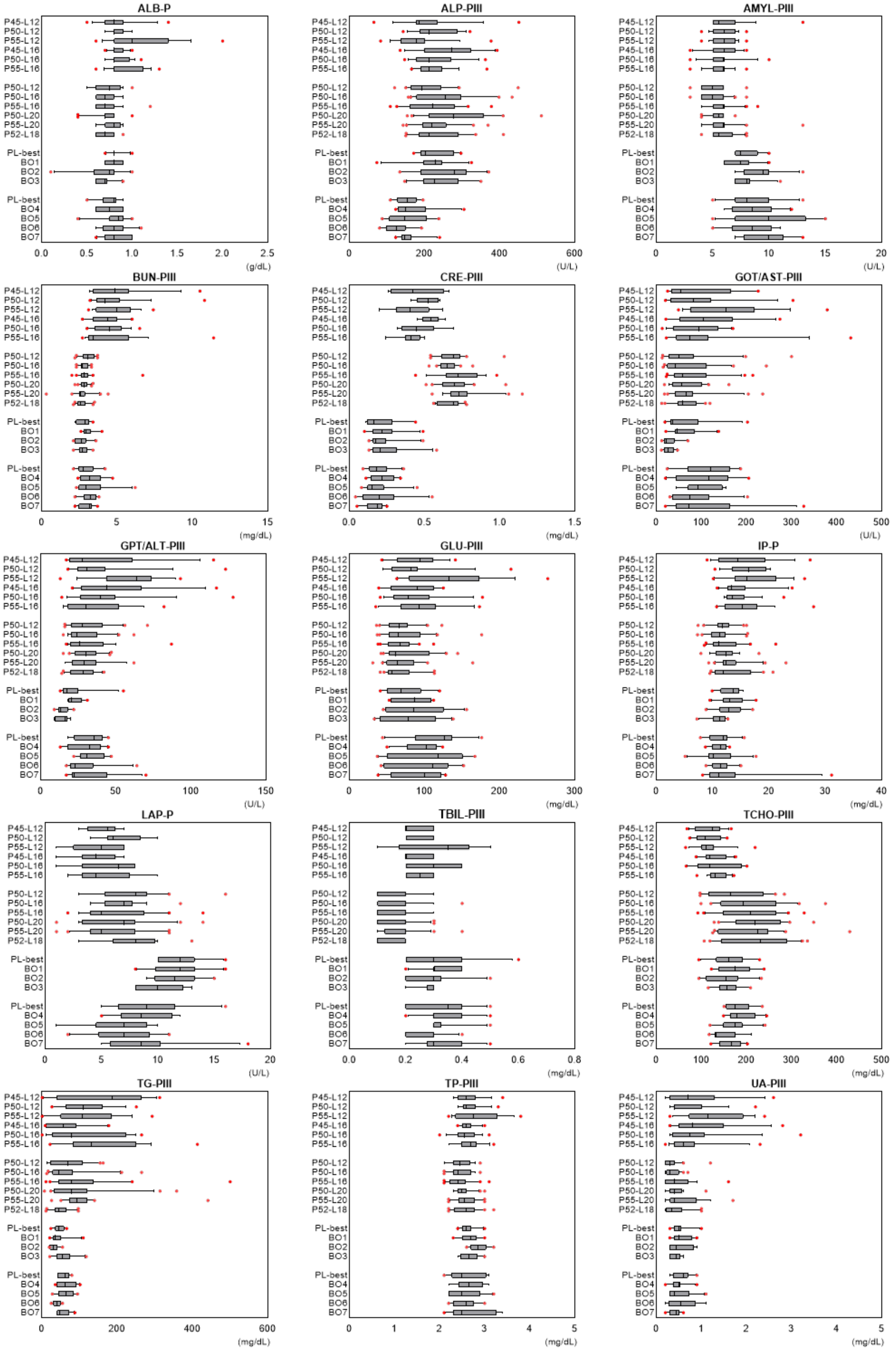
ALB-P
ALP-PIII
AMYL-PIII
BUN-PIII
CRE-PIII
GOT/AST-PIII
GPT/ALT-PIII
GLU-PIII
IP-P
LAP-P
TBIL-PIII
TCHO-PIII
TG-PIII
TP-PIII
UA-PIII
P45-L12
P50-L12
P55-L12
P45-L16
P50-L16
P55-L16
P50-L12
P50-L16
P55-L16
P50-L20
P55-L20
P52-L18
PL-best
BO1
BO2
BO3
PL-best
BO4
BO5
BO6
BO7
0.0 0.5 1.0 1.5 2.0 2.5
(g/dL)
0 200 400 600
(U/L)
0 5 10 15 20
(U/L)
0 5 10 15
(mg/dL)
0.0 0.5 1.0 1.5
(mg/dL)
0 100 200 300 400 500
(U/L)
0 50 100 150
(U/L)
0 100 200 300
(mg/dL)
0 10 20 30 40
(mg/dL)
0 5 10 15 20
(U/L)
0.0 0.2 0.4 0.6 0.8
(mg/dL)
0 100 200 300 400 500
(mg/dL)
0 200 400 600
(mg/dL)
0 1 2 3 4 5
(mg/dL)
0 1 2 3 4 5
(mg/dL)

Figure S2. The results of biochmical analysis are expressed by box-and-whisker plots with 10 to 90 percentiles. The black bar in the plots indicates the median. Alanine aminotransferase (GPT/ALT-PⅢ), alkaline phosphatase (ALP-PⅢ), amylase (AMYL-PⅢ), aspartate aminotransferase (GOT/AST-PⅢ), inorganic phosphorus (IP-P), leucine aminopeptidase (LAP-P), albumin (ALB-PⅢ), creatinine (CRE-PⅢ), glucose (GLU-PⅢ), total cholesterol(TCHO-PⅢ), total protein (TP-PⅢ), total bilirubin (TBIL-PⅢ), and triglyceride(TG-PⅢ), urea-nitrogen (BUN-PⅢ), and uric acid (UA-PⅢ).

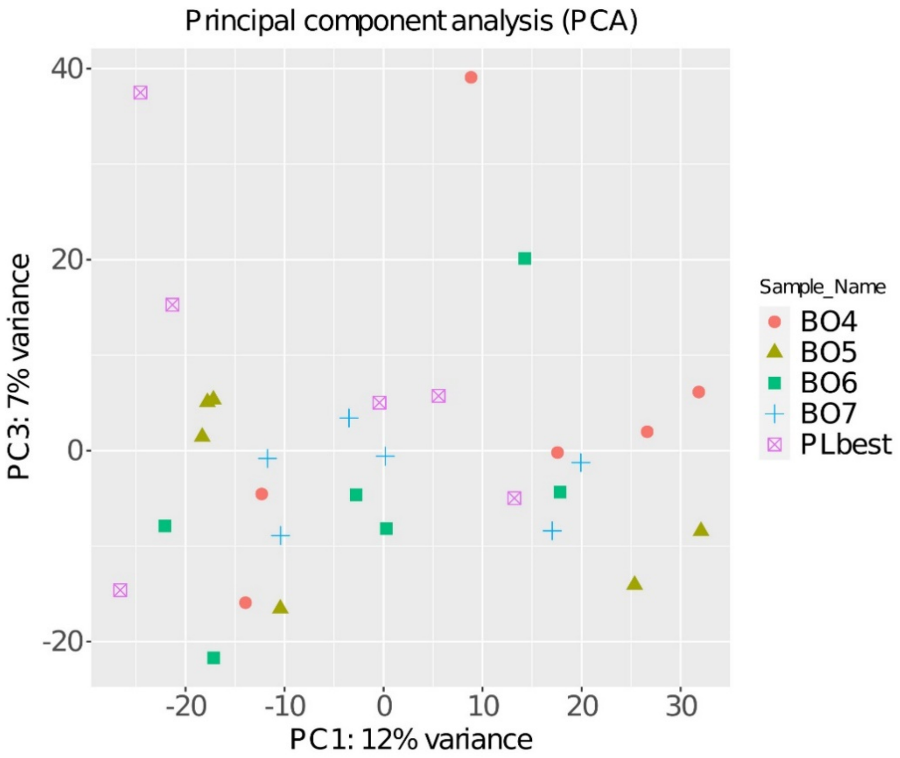

Figure S3. The cauterization of the experimental groups by PCA analysis. The graph was constructed using PC1 and PC3. Red circle:BO4 group, yellow triangle: BO5, green square: BO6, blue plus symbol: BO7, purple square cross symbol: PL-best.

Table S4. Formulation and proximate composition of the test diets for R2-1. The detailed compositions of the Vitamin and Mineral mixtures are described in Table S1.

| | Control | Trp 0.5 | Trp 1 | Trp 2 |
|---|---|---|---|---|
| Ingredients (g/kg) | | | | |
| Fish meal (Anchovy/Sardine) | 510 | 505 | 500 | 490 |
| Krill meal | 100 | 100 | 100 | 100 |
| Wheat flour | 228 | 228 | 228 | 228 |
| Tapioca starch | 90 | 90 | 90 | 90 |
| Fish oil | 15 | 15 | 15 | 15 |
| Vitamin mixture | 27 | 27 | 27 | 27 |
| Mineral mixture | 30 | 30 | 30 | 30 |
| L-Tryptophan (Trp) | 0 | 5 | 10 | 20 |
| Proximate composition (% dry matter basis) | | | | |
| Moisture | 5.5 | 5.0 | 5.0 | 6.3 |
| Crude protein (%) | 48.5 | 47.9 | 48.2 | 47.4 |
| Crude Lipid (%) | 10.8 | 10.3 | 10.6 | 10.3 |
| Ash | 10.5 | 104. | 10.4 | 10.1 |

Table S5. Formulation and proximate composition of the test diets for S2-2. The detailed compositions of the Vitamin and Mineral mixtures are described in Table S1.

| R3-1 | P50-L12 (Control) | IMP 0.25% | IMP 0.5% | GAA 0.1% | GAA 0.2% | Cre 0.6% |
|---|---|---|---|---|---|---|
| Ingredients (g/kg) | | | | | | |
| Fish meal (Anchovy/Sardine) | 530 | 530 | 530 | 530 | 530 | 530 |
| Krill meal | 110 | 120 | 110 | 110 | 110 | 120 |
| Wheat flour | 173 | 173 | 173 | 173 | 173 | 173 |
| Tapioca starch | 80 | 80 | 80 | 80 | 80 | 80 |
| Fish oil | 50 | 50 | 50 | 50 | 50 | 50 |
| Vitamin mixture[1] | 27 | 27 | 27 | 27 | 27 | 27 |
| Mineral mixture[1] | 30 | 30 | 30 | 30 | 30 | 30 |
| Inosinic acid (IMP) | 0 | 2.5 | 5 | 0 | 0 | 0 |
| Guanidinoacetate (GAA) | 0 | 0 | 0 | 1 | 2 | 0 |
| Creatine (Cre) | 0 | 0 | 0 | 0 | 0 | 6 |
| Proximate composition (% dry matter basis) | | | | | | |
| Moisture | 6.7 | 7.2 | 5.6 | 5.6 | 5.4 | 8.0 |
| Crude protein (%) | 50.2 | 50.4 | 50.9 | 50.2 | 50.5 | 51.3 |
| Crude Lipid (%) | 14.5 | 14.4 | 14.3 | 14.3 | 14.3 | 14.3 |
| Ash | 12.5 | 12.6 | 12.6 | 12.4 | 12.4 | 12.4 |

Table S6. Formulation and proximate composition of the test diets for S2-3. The detailed compositions of the Vitamin and Mineral mixtures are described in Table S1.

| R3-1 | PL-best | Gly 1% | Gly 2% | Lac 1% | Lac 2% | Cre 0.3% |
|---|---|---|---|---|---|---|
| Ingredients (g/kg) | | | | | | |
| Fish meal (Anchovy/Sardine) | 570 | 570 | 570 | 570 | 570 | 570 |
| Krill meal | 120 | 120 | 120 | 120 | 120 | 120 |
| Wheat flour | 123 | 123 | 123 | 123 | 123 | 123 |
| Tapioca starch | 40 | 40 | 40 | 40 | 40 | 40 |
| Fish oil | 90 | 90 | 90 | 90 | 90 | 90 |
| Vitamin mixture | 27 | 27 | 27 | 27 | 27 | 27 |
| Mineral mixture | 30 | 30 | 30 | 30 | 30 | 30 |
| Glycine (Gly) | 0 | 10 | 20 | 0 | 0 | 0 |
| Lactic acid (Lac) | 0 | 0 | 0 | 10 | 20 | 0 |
| Creatine (Cre) | 0 | 0 | 0 | 0 | 0 | 3 |
| Proximate composition (% dry matter basis) | | | | | | |
| Moisture | 5.4 | 6.5 | 5.4 | 5.7 | 5.5 | 6.3 |
| Crude protein (%) | 53.4 | 54.4 | 55.0 | 52.1 | 52.1 | 54.0 |
| Crude Lipid (%) | 18.8 | 18.8 | 18.1 | 18.8 | 18.0 | 18.2 |
| Ash | 13.9 | 13.9 | 13.5 | 14.3 | 15.0 | 13.8 |

Table S7. Formulation and proximate composition of the test diets for S2-4. The detailed compositions of the Vitamin and Mineral mixtures are described in Table S1.

| R3-1 | PL-best | Cys 1% | Cys 2% | Arg 1.5% | Arg 3% | Cho 0.25% |
|---|---|---|---|---|---|---|
| Ingredients (g/kg) | | | | | | |
| Fish meal (Anchovy/Sardine) | 570 | 570 | 570 | 570 | 570 | 570 |
| Krill meal | 120 | 120 | 120 | 120 | 120 | 120 |
| Wheat flour | 123 | 123 | 123 | 123 | 123 | 123 |
| Tapioca starch | 40 | 40 | 40 | 40 | 40 | 40 |
| Fish oil | 90 | 90 | 90 | 90 | 90 | 90 |
| Vitamin mixture1 | 27 | 27 | 27 | 27 | 27 | 27 |
| Mineral mixture2 | 30 | 30 | 30 | 30 | 30 | 30 |
| Cysteine (Cys) | 0 | 10 | 20 | 0 | 0 | 0 |
| Arginine (Arg) | 0 | 0 | 0 | 15 | 30 | 0 |
| Choline (Cho) | 0 | 0 | 0 | 0 | 0 | 5 |
| Proximate composition (% dry matter basis) | | | | | | |
| Moisture | 6.1 | 7.4 | 7.9 | 7.1 | 9.1 | 6.0 |
| Crude protein (%) | 53.8 | 54.0 | 53.5 | 55.4 | 57.3 | 53.7 |
| Crude Lipid (%) | 18.6 | 18.6 | 18.4 | 18.8 | 18.2 | 17.9 |
| Ash | 14.0 | 13.7 | 13.6 | 13.5 | 13.4 | 13.9 |

Table S8. First additive recommendation by BO. Asterisked items were used in the feeding experiments at S2-5.

| Optimization of feeding efficiency (FE) | | | | | | |
|---|---|---|---|---|---|---|
| | Gly | Trp | IMP | Cho | Scaled FE (pred.) | Variance of scaled FE (pred.) |
| 1 | 0 | 0 | 1 | 0 | 1.05 | 0.37 |
| 2 | 2 | 0 | 0.75 | 0 | 1.05 | 0.37 |
| 3* (BO1) | 0 | 0 | 0.75 | 0.25 | 1.05 | 0.37 |
| Optimization of dairy growth rates (DGR) | | | | | | |
| | Gly | Trp | IMP | Cho | Scaled DGR (pred.) | Variance of scaled DGR (pred.) |
| 1* (BO3) | 1 | 0.5 | 0.75 | 0 | 1.06 | 0.36 |
| 2* (BO1) | 0 | 0 | 0.75 | 0.25 | 1.05 | 0.37 |
| 3 | 0 | 0 | 1 | 0 | 1.05 | 0.36 |
| Optimization of the multiple of FE and DGR | | | | | | |
| | Gly | Trp | IMP | Cho | Multiple of scaled FE and DGR (pred.) | Variance of multiple of scaled FE and DGR (pred.) |
| 1* (BO2) | 1 | 0 | 0.75 | 0 | 1.08 | 0.35 |
| 2* (BO3) | 0 | 0.5 | 0.75 | 0 | 1.09 | 0.33 |
| 3 | 0 | 0 | 0.5 | 0.25 | 1.05 | 0.36 |

Table S9. Formulation and proximate composition of the test diets for S2-5. The detailed compositions of the Vitamin and Mineral mixtures are described in Table S1.

| R3-1 | Control (P51-L18) | Cho-0.5 | BO1 | BO2 | BO3 |
|---|---|---|---|---|---|
| Ingredients (g/kg) | | | | | |
| Fish meal (Anchovy/Sardine) | 570 | 570 | 570 | 570 | 565 |
| Krill meal | 120 | 120 | 120 | 120 | 120 |
| Wheat flour | 123 | 123 | 123 | 123 | 123 |
| Tapioca starch | 40 | 40 | 40 | 40 | 40 |
| Fish oil | 90 | 90 | 90 | 90 | 90 |
| Vitamin mixture | 20 | 20 | 20 | 20 | 20 |
| Mineral mixture | 30 | 30 | 30 | 30 | 30 |
| L-Tryptophan (Trp) | 0 | 0 | 0 | 0 | 5 |
| Glycine (Gly) | 0 | 0 | 0 | 10 | 0 |
| Inosinic acid (IMP) | 0 | 0 | 7.5 | 7.5 | 7.5 |
| Choline (Cho) | 0 | 10 | 5 | 0 | 0 |
| Proximate composition (% dry matter basis) | | | | | |
| Moisture | 4.3 | 4.6 | 5.2 | 4.6 | 6.3 |
| Crude protein (%) | 51.3 | 51.9 | 51.7 | 52.0 | 51.5 |
| Crude Lipid (%) | 18.6 | 18.6 | 18.4 | 18.8 | 18.2 |
| Ash | 15.2 | 13.8 | 14.1 | 14.0 | 14.0 |

Table S10. Second additive recommendation by BO

Optimization of feeding efficiency (FE)

| | Gly | Trp | IMP | Cho | Scaled FE (pred.) | Variance of scaled FE (pred.) |
|---|---|---|---|---|---|---|
| 1* (BO4) | 0 | 0.5 | 0 | 0.25 | 1.03 | 0.22 |
| 2 | 1 | 0.5 | 0 | 0 | 1.02 | 0.21 |
| 3* (BO7) | 1 | 0 | 0.5 | 0.25 | 0.97 | 0.24 |

Optimization of Dairy Growth Rates (DGR)

| | Gly | Trp | IMP | Cho | Scaled DGR (pred.) | Variance of scaled DGR (pred.) |
|---|---|---|---|---|---|---|
| 1* (BO4) | 0 | 0.5 | 0 | 0.25 | 1.01 | 0.23 |
| 2 | 1 | 0.25 | 0.25 | 0.25 | 0.93 | 0.29 |
| 3* (BO6) | 2 | 0.5 | 0.5 | 0 | 0.87 | 0.33 |

Optimization of the multiple of FE and DGR

| | Gly | Trp | IMP | Cho | multiple of FE and DGR (pred.) | Variance of multiple of FE and DGR (pred.) |
|---|---|---|---|---|---|---|
| 1* (BO4) | 0 | 0.5 | 0 | 0.25 | 1.09 | 0.23 |
| 2* (BO5) | 1 | 0.5 | 0 | 0.25 | 0.94 | 0.3 |
| 3* (BO6) | 2 | 0.5 | 0.5 | 0 | 0.88 | 0.33 |

Table S11. Formulation and proximate composition of the test diets for R4-2. The detailed compositions of the Vitamin and Mineral mixtures are described in Table S1.

| R3-1 | Control (P52-L18) | BO4 (Multi1) | BO5 (Multi2) | BO6 (Multi3) | BO7 (FE 3) |
|---|---|---|---|---|---|
| Ingredients (g/kg) | | | | | |
| Fish meal (Anchovy/Sardine) | 570 | 565 | 565 | 565 | 565 |
| Krill meal | 120 | 120 | 120 | 120 | 120 |
| Wheat flour | 123 | 123 | 123 | 123 | 123 |
| Tapioca starch | 40 | 40 | 40 | 40 | 40 |
| Fish oil | 90 | 90 | 90 | 90 | 90 |
| Vitamin mixture | 20 | 20 | 20 | 20 | 20 |
| Mineral mixture | 30 | 30 | 30 | 30 | 30 |
| L-Tryptophan (Trp) | 0 | 5 | 5 | 5 | 5 |
| Glycine (Gly) | 0 | 0 | 10 | 20 | 10 |
| Inosinic acid (IMP) | 0 | 0 | 0 | 5 | 5 |
| Choline (Cho) | 0 | 5 | 5 | 0 | 5 |
| Proximate composition (% dry matter basis) | | | | | |
| Moisture | 7.7 | 6.6 | 7.2 | 8.4 | 7.5 |
| Crude protein (%) | 51.8 | 53.0 | 53.7 | 53.8 | 53.3 |
| Crude Lipid (%) | 18.4 | 18.6 | 18.7 | 18.5 | 17.9 |
| Ash | 12.1 | 12.5 | 12.2 | 12.2 | 12.0 |
